# Inference of Species Phylogenies from Bi-allelic Markers Using Pseudo-likelihood

**DOI:** 10.1101/289207

**Authors:** Jiafan Zhu, Luay Nakhleh

## Abstract

**Motivation:** Phylogenetic networks represent reticulate evolutionary histories. Statistical methods for their inference under the multispecies coalescent have recently been developed. A particularly powerful approach uses data that consist of bi-allelic markers (e.g., single nucleotide polymorphism data) and allows for exact likelihood computations of phylogenetic networks while numerically integrating over all possible gene trees per marker. While the approach has good accuracy in terms of estimating the network and its parameters, likelihood computations remain a major computational bottleneck and limit the method’s applicability.

**Results:** In this paper, we first demonstrate why likelihood computations of networks take orders of magnitude more time when compared to trees. We then propose an approach for inference of phylogenetic networks based on pseudo-likelihood using bi-allelic markers. We demonstrate the scalability and accuracy of phylogenetic network inference via pseudo-likelihood computations on simulated data. Furthermore, we demonstrate aspects of robustness of the method to violations in the underlying assumptions of the employed statistical model. Finally, we demonstrate the application of the method to biological data. The proposed method allows for analyzing larger data sets in terms of the numbers of taxa and reticulation events. While pseudo-likelihood had been proposed before for data consisting of gene trees, the work here uses sequence data directly, offering several advantages as we discuss.

**Availability:** The methods have been implemented in PhyloNet (http://bioinfocs.rice.edu/phylonet).

## 1 Introduction

Species phylogenies model how species evolve and diversify. When species split and diversify without subsequent exchange of genetic material between different species, the species phylogeny takes the shape of a tree. In the post-genomic era, the inference of species trees in general makes use of the availability of sequence data of multiple individual loci across the genomes. Methods for species tree inference from such data view evolution of the sequence data as two co-occurring stochastic processes: one that models the growth of the genealogies of individual loci within the branches of the species tree and another that models the evolution of sequences of individual loci along the branches of the corresponding genealogies. The latter process is most commonly captured by Markov models of sequence evolution and is the basis for computing the likelihood of (gene) trees [6]. Modeling the former process depends on the evolutionary processes acting on the individual loci. In particular, the coalescent model [9] is one such model of growth of the genealogies when individual loci evolve under the idealized Wright-Fisher model. When the species tree is viewed as a set of populations stitched together according to the tree structure, the coalescent model is extended into the multispecies coalescent [5].

However, when different species exchange genetic material, the tree structure is no longer adequate for modeling their resulting reticulate evolutionary history. It is now well established that the evolutionary histories of several groups of species, both prokaryotic and eukaryotic, are reticulate [13]. In prokaryotes, the most common process of reticulation is horizontal gene transfer [8, 10], whereas hybridization and introgression are the main processes of reticulation in eukaryotes [1, 2, 19, 11, 12, 18, 7, 16, 13, 27]. A phylogenetic network is a rooted, directed acyclic graphs that represents the evolutionary histories of a set of taxa when that evolutionary history is not strictly treelike [15].

To extend the aforementioned model of two co-occurring stochastic processes to reticulate evolutionary histories, the multispecies network coalescent (MSNC) was devised [33, 34]. Under this model, the genealogy of an individual locus evolves within the branches of a phylogenetic network, thus capturing evolutionary scenarios where loci could be exchanged across species boundaries. The model enabled the development of statistical methods for inferring phylogenetic networks from multi-locus data sets, summarized in Table 1.

**Table 1:** Methods for phylogenetic network inference under the MSNC. 𝓛: likelihood computation (F: full; P: pseudo); GT: gene trees; GTT: gene tree topologies; SA: sequence alignments; BM: bi-allelic markers.

| Data | $\mathcal{L}$ | Reference |
| --- | --- | --- |
| GT/GTT | F | [34, 28] |
| GTT | P | [31, 21] |
| SA | F | [26, 35] |
| BM | F | [36] |
| BM | P | This paper |

In terms of the data that they utilize, statistical methods for phylogenetic network inference based on the multispecies network coalescent can be grouped in three categories. The methods of [33], [34], and [28] use gene tree estimates as the input data and infer the phylogenetic network and its parameters based on maximum likelihood or Bayesian inference. A major computational bottleneck for these methods is the calculation of the likelihood of phylogenetic networks. These calculations can take hours and even days for a single network if the numbers of taxa and reticulations are large. This was the main motivation behind the development of pseudo-likelihood methods for phylogenetic networks [31, 21]. However, two disadvantages of using gene tree topology estimates as the input data (for both the full- and pseudo-likelihood methods) are the inability to estimate some parameters of interest that could be estimable from sequence data, and in fact the gene tree estimates could have much error in them, especially at the scale of evolution where incomplete lineage sorting is a concern. To address these two issues, the methods of [26], and [35] employ Bayesian Markov chain Monte Carlo to sample the posterior of phylogenetic networks and gene trees from the sequence data directly, where the data for each locus consist of a sequence alignment. Most recently, [36] devised a Bayesian method for inferring phylogenetic networks from data that consist of unlinked bi-allelic markers (such as single nucleotide polymorphism data and amplified fragment length polymorphisms).

While the method of [36] is very promising in terms of accuracy and has the advantage that it performs numerical integration that completely sidesteps the issue of sampling the enormous gene tree space, it is very slow in practice, rendering the inference of networks with more than 5-6 taxa and 3-4 reticulations infeasible. This paper concerns the scalability of the method. We first discuss factors that make likelihood computations on networks computationally orders of magnitudes more demanding that on trees even when the network and tree differ by a single reticulation. We then propose a pseudo-likelihood formulation based on bi-allelic markers, along with an inference method. We demonstrate the accuracy, robustness, and speed of the method and its ability to analyze much larger data sets than full-likelihood-based methods can handle. The developed methods are implemented in the open-source, publicly available software package PhyloNet [23, 29].

## 2 Background

A *phylogenetic network* Ψ on set 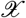 of taxa is a rooted, directed, acyclic graph whose leaves are bijectively labeled by 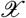. Each node in the network has in-degree of 0 (the root), 1 (a tree node), or 2 (a reticulation node). The out-degree of each node is at most 2. Network Ψ’s sets of nodes and edges are denoted by *V* (Ψ) and *E*(Ψ), respectively.

Each node in the network has a species divergence time and each edge *b* has an associated population mutation rate *θ_b_* = 4*N_b_µ* where *N*_*b*_ is the effective population size associated with edge *b* and *µ* is the mutate rate per site per generation. For calculations under the coalescent, it is always assumed that there is an infinite-length edge above the root of the network. Furthermore, for every pair of reticulation edges *e*_1_ and *e*_2_ that share the same reticulation node, we associate an inheritance probability, *γ*, such that 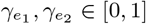 with 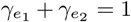. We denote by Γ the vector of inheritance probabilities corresponding to all the reticulation nodes in the phylogenetic network. In order to simplify notation, we assume here that Ψ represents the phylogenetic network topology and all the other parameters; that is, the divergence times, population mutation rates, and inheritance probabilities are components of Ψ, in addition to the topology.

In this paper we assume the multispecies network coalescent process. Consider a data set 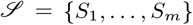 where *S*_*i*_ consists of the binary states of *m* unlinked (independent) bi-allelic markers for a set 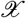 of taxa. The likelihood of a species phylogeny Ψ (topology and parameters) is given by

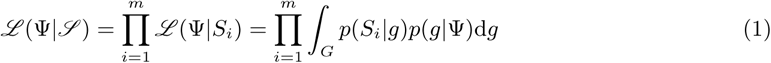

where the integration is taken over all possible gene trees. The term *p*(*S_i_|g*) is the likelihood of gene tree *g* given the sequence data of locus *i* [6] and *p*(*g|*Ψ) is the density function of gene trees given the species phylogeny and its parameters [17].

[3] introduced an algorithm for analytically computing the integration in Eq. (1) for bi-allelic markers, thus avoiding the need to sample gene trees to estimate the integral. [36] extended the method of Bryant *et al.* in novel ways so that the integration in Eq. (1) can be done analytically also when the species phylogeny is a network.

## 3 Methods

We first discuss factors that govern the computational complexity of full likelihood calculations on networks, and then propose a pseudo-likelihood function of phylogenetic networks and demonstrate its scalability.

### 3.1 When it comes to computational complexity, a network is not merely a tree with a few additional reticulations

While the full likelihood computations of the algorithm of [3] allowed for inferring species trees with tens of taxa, the computational complexity of these computations exploded when the species phylogeny was a network. We now explain the explosion in the running time of computing the probability of a gene tree topology on a network as compared to that of computing the probability of a gene tree on the species tree “inside” the network. While the computations in this paper do not involve gene tree probability computations, the limitations of the likelihood computations of [36] follow the same rationale.

The probability mass function (pmf) *p*(*g|*Ψ), where *g* is a gene tree topology, is central to statistical inference of species trees and networks. In the case of species trees, Ψ is a tree, and Γ is irrelevant. As was shown in [4, 34], 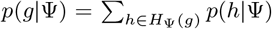, where *H*_Ψ_(*g*) is the set of all coalescent histories of *g* inside Ψ. Roughly speaking, a coalescent history of *g* is an embedding of *g* within the branches of Ψ.

The size of *H*_Ψ_(*g*) for a gene tree topology *g* when Ψ is a tree is exponential in the number of leaves in the gene tree [20, 22]. [30] devised an efficient way of computing the pmf for species trees without explicitly enumerating all coalescent histories in the set *H*_Ψ_(*g*). Similarly, [32] devised an efficient way of computing the pmf for species networks without enumerating all coalescent histories. However, with this improved method, likelihood calculations for species trees scale very well, whereas that is not the case for phylogenetic networks.

Here, we extended the algorithm of [22] to compute the size of *H*_Ψ_(*g*) for a gene tree topology *g* and phylogenetic network Ψ. For simplicity, we assume one individual is sampled per species so that the leaves of the gene tree and phylogenetic network are labeled by the same set 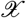 of taxa (the algorithm can be trivially extended to the case where multiple individuals are sampled per species). Every edge *e* in a gene tree defines a *cluster*, *c*_*e*_, which is the set of leaves under the edge. We denote by *L*(*t*) the set of all taxa labeling the leaves of tree *t*. Let *C*_*g*_ be the set of all clusters of size *≥* 2 of taxa in the gene tree *g*, and let *X*_*g*_ = *L*(*g*). Assume *E*(Ψ) includes a special edge *r* that is incoming into the root. We write (*x, y*) to denote a tree whose root has two children that are subtrees *x* and *y*. We define function *ρ*_*c*_(*e*) to represent the number of coalescent scenarios of leaves in set *c* ∩ *c*_*e*_. We define *children*(*e*) for edge *e* = (*u, v*) *∈ E*(Ψ) as the set of all edges (*v, x*) *∈ E*(Ψ) for *x ∈ V* (Ψ). For two edges *e*_1_, *e*_2_*∈ E*(Ψ), *e*_1_ ≠ *e*_2_, we say that *e*_2_ = (*u*_2_, *v*_2_) is on the path from *e*_1_ = (*u*_1_, *v*_1_) to the root, denoted by *e*_1_ ≺ *e*_2_, if there is path from the root of Ψ to *u*_1_ that passes through both *u*_2_ and *v*_2_. If no such path exists, we write *e*_1_⊀ *e*_2_. Denoting by *ζ*(*e*_1_, *e*_2_) the number of paths from *e*_1_ to *e*_2_, the quantity can be computed by:

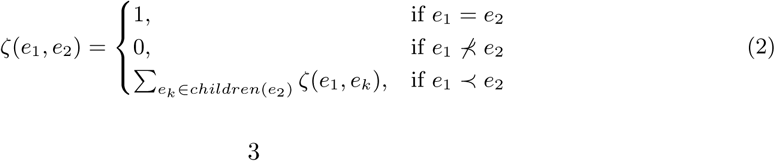

Using the function *ζ*, *ρ_c_*(*e*) is computed using Algorithm 1. The size of *H*_Ψ_(*g*) is given by 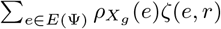.

#### Algorithm 1: Compute *ρ_c_*(*e*)

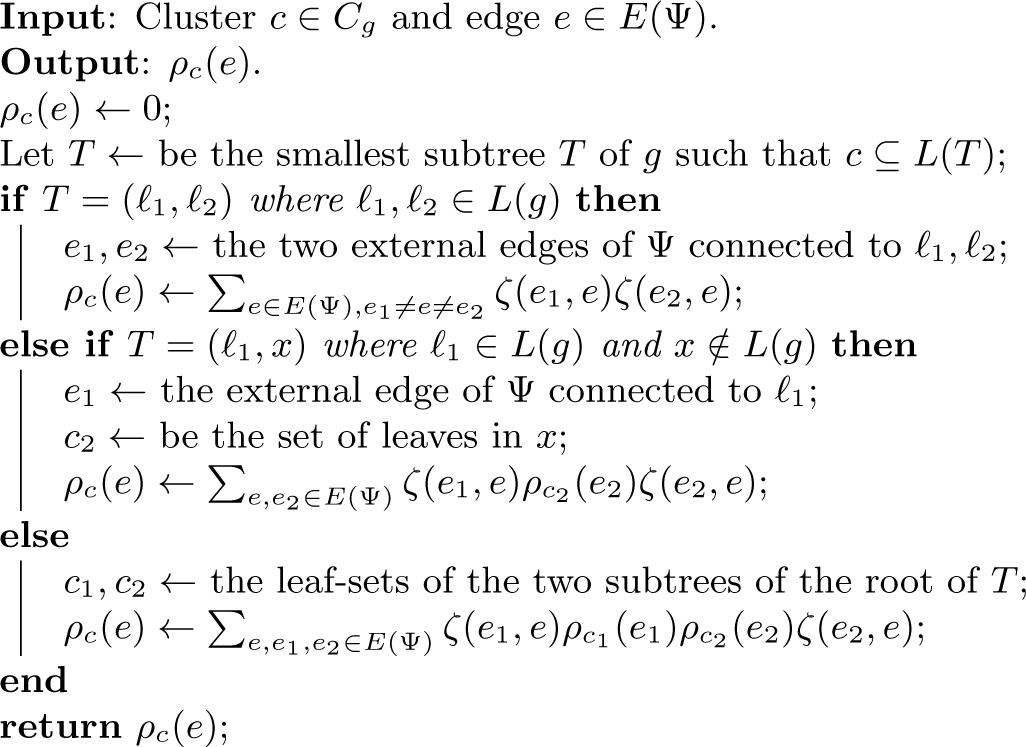

Fig. 1A demonstrates the effect on the size of set *H* of the addition of a single reticulation to an underlying tree to form a network. For some networks, going from a tree to a network with a single reticulation increased the size of *H* by ten million fold. The efficient algorithms of [30], and [32] ameliorate the computational complexity of computing the pmf by employing a bottom-up algorithm that stores values at nodes. In other words, they trade off memory for time. The reason that these algorithms scale in the case of trees but not networks is precisely the illustration in Fig. 1A.

**Figure 1:**
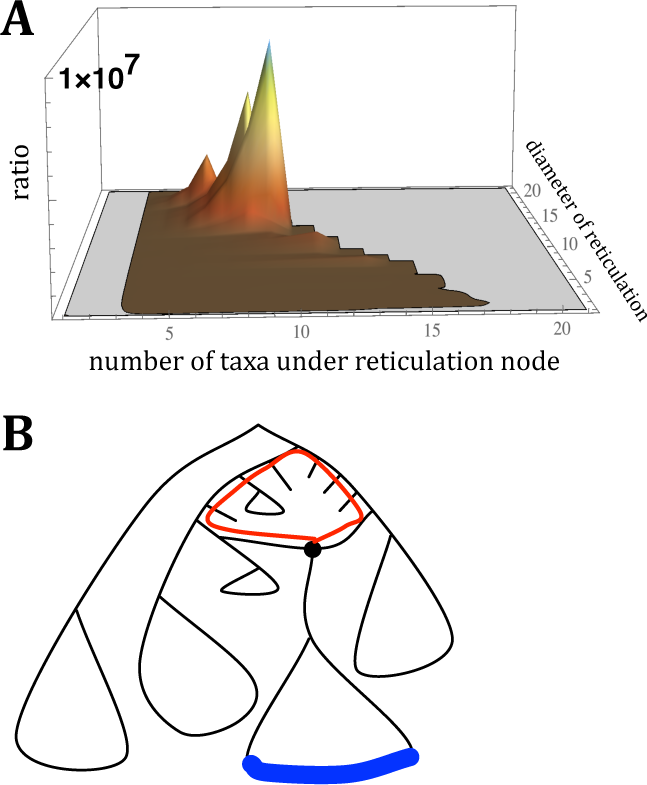
The ratio of the number of coalescent histories on a network to the number of coalescent histories on the underlying tree. (**A**) The results are based on 30 random 20-leaf phylogenetic networks and 100 gene trees for each network. Each of the 30 networks was obtained by adding a single reticulation to an underlying tree. (**B**) A phylogenetic network with a single reticulation. The diameter of the reticulation is the number of edges on the paths marked with the red cycle. The taxa under the reticulation correspond to the leaves marked with the blue line.

An obvious factor that significantly affects the running time of computing the pmf is the number of taxa (leaves) in the network. However, in some cases, computing the likelihood of a network on 20 taxa could take less time than that of computing the likelihood of a network with, say, 10 taxa. The complexity of a phylogenetic network is governed by the diameters of the reticulation nodes and the number of leaves under the reticulation nodes (Fig. 1B). The larger either or both of these, the worse the explosion in the size of *H* is and, consequently, the worse the likelihood calculations become in terms of time and memory requirements. These exorbitant computational costs are only exacerbated, in a potentially exponential manner, when the number of reticulations increases.

### 3.2 Pseudo-likelihood

Given a phylogenetic network Ψ on set 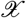 of taxa, a subset Ψ′ of Ψ on subset 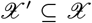 of taxa is the phylogenetic network obtained by restricting Ψ to the leaves in 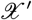 (if there are multiple paths between two leaves, we keep them all). We denote such a subnet by 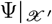. Fig. 2 shows a phylogenetic network and its four three-taxon subnets. It is important to note nodes and reticulation edges in the subnets have the same divergence times and inheritance probabilities as their corresponding nodes and reticulation edges, respectively, in the full phylogenetic network. We assume the same population size across all branches of the network.

**Figure 2:**
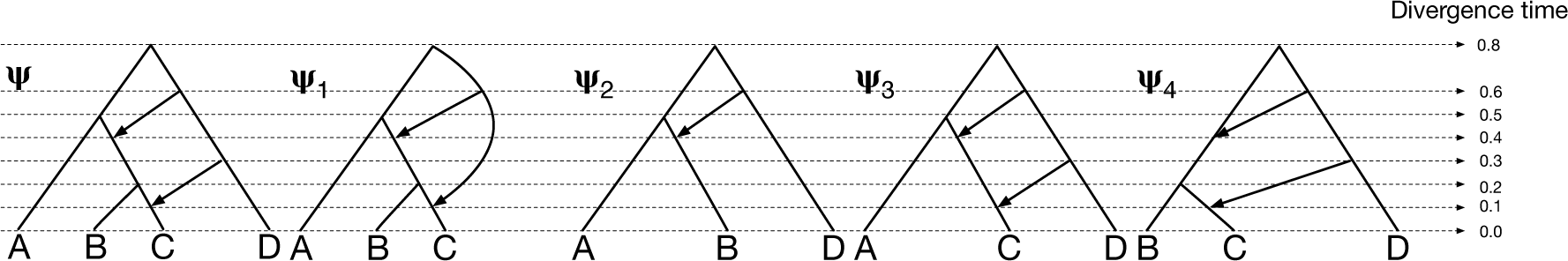
A phylogenetic network and its 3-taxon subnets. Phylogenetic network Ψ induces four subnets Ψ_1_, Ψ_2_, Ψ_3_, and Ψ_4_. The inheritance probabilities are the same among the network and subnets for corresponding reticulation edges, and the times of all corresponding nodes are the same among the network and its three subnets.

While subnets with 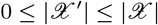 could be considered, here we consider only three-taxon subnets. For a phylogenetic network Ψ, we denote by 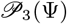 the set of all three-taxon subnets of Ψ, also called trinets.

Let Ψ be a (parameterized) phylogenetic network on set 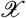 of taxa, and let 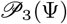 be the set of Ψ’s trinets. Consider a data set 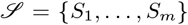 where *S*_*i*_ consisting of the binary states of *m* unlinked (independent) bi-allelic markers for a set 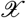 of taxa. We denote by 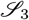 the set of all data sets obtained by restricting 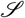 to all three-taxon subsets. Table 2 shows an example of 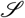 and 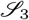 for a four-taxon phylogenetic network (e.g., the one in Fig. 2). The pseudo-likelihood of a species phylogeny Ψ (topology and parameters) is given by

**Table 2:** Site patterns on the phylogenetic network Ψ of Fig. 2 and its marginalized patterns on the four subnets.

| | $\mathcal{S}(\Psi)$ | | | $\mathcal{S}_3(\Psi_1)$ | | | $\mathcal{S}_3(\Psi_2)$ | | | $\mathcal{S}_3(\Psi_3)$ | | | $\mathcal{S}_3(\Psi_4)$ | | |
| --- | --- | --- | --- | --- | --- | --- | --- | --- | --- | --- | --- | --- | --- | --- | --- |
|  | s1 | s2 | s3 | s1 | s2 | s3 | s1 | s2 | s3 | s1 | s2 | s3 | s1 | s2 | s3 |
| A | 1 | 0 | 0 | 1 | 0 | 0 | 1 | 0 | 0 | 1 | 0 | 0 |  |  |  |
| B | 0 | 1 | 0 | 0 | 1 | 0 | 0 | 1 | 0 |  |  |  | 0 | 1 | 0 |
| C | 1 | 1 | 1 | 1 | 1 | 1 |  |  |  | 1 | 1 | 1 | 1 | 1 | 1 |
| D | 1 | 1 | 1 |  |  |  | 1 | 1 | 1 | 1 | 1 | 1 | 1 | 1 | 1 |

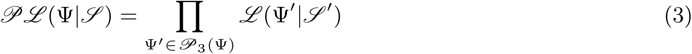

where 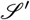 is the data set obtained by restricting 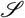 to the taxa in Ψ′, and 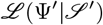 is computed according to Eq. (1) using the algorithm of [36]. It is important to note here that while the exposition is given in terms of phylogenetic networks and the algorithm of [36], this same formulation applies to species trees and the algorithm of [3].

#### Searching the phylogenetic network space

We use simulated annealing to search the phylogenetic network space with similar moves and setting to that of [31], with two main differences. First, when a delete-reticulation move is proposed, it is accepted with probability 0.01 no matter how the pseudo-likelihood changes. In our testing of the search strategy, we found that such a modification helps the search jump out of locally optimal states. Second, the optimal network with one fewer reticulation nodes than the the maximum number was kept during each iteration of search, and it was used as the starting state for every subsequent iteration except the first one. The search is conducted in a number of iterations, where each iteration walks the space of phylogenetic networks starting at some temperature that is reduced gradually during the iteration. So, for example, an iteration could inspect on the order of 50,000 points in the parameter space.

## 4 Results

### 4.1 Simulations

We first set out to assess the running time of computing the pseudo-likelihood of phylogenetic networks of varying sizes. To achieve this, we generated networks with 10, 20, 50, and 100 taxa and with 0, 1, 2, 3, and 4 reticulations. We used PhyloGen [24] to first generate random species trees with 10, 20, 50, and 100 taxa. Then, for each species tree, we randomly added 0, 1, 2, 3, and 4 reticulations. To add a reticulation to a species network, we selected two edges uniformly at random and added an edge between their midpoints in a direction that ensures no cycles are created. Inheritance probabilities were assigned 0.5 to reticulation edges. Then, we used PhyloNet [29] to simulate the evolution of 10000 bi-allelic markers on each species network. Finally, we computed the pseudo-likelihoods of each phylogenetic network using the data sets generated on it. The results are given in Fig. 3.

**Figure 3:**
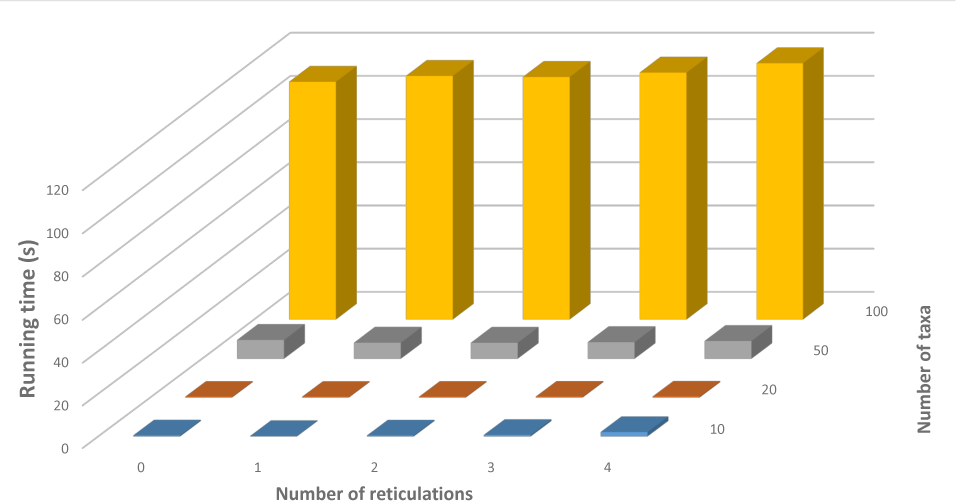
Running times of computing the pseudo-likelihood of networks of varying numbers of taxa and leaves. The running times are reported in seconds. They were measured on a desktop computer with 16G RAM and INTEL XEON E3-1245 @ 3.5GHz, and 8 threads were used.

As the results demonstrate, computing the pseudo-likelihood is very fast. In particular, it is worth pointing out that computing the full likelihood of networks of with more than 10 taxa and 3 or 4 reticulations could be impractical for many topologies. As discussed above, this is governed not only by the number of taxa and number of reticulation nodes, but more importantly by the diameters of the reticulation nodes, the dependence among the cycles in the underlying undirected graph of the network, and the numbers of taxa that are descendants of reticulation nodes. The pseudo-likelihood calculations take fraction of a second on phylogenetic networks with 20 or fewer taxa, regardless of the number of reticulations. The calculations take about 10 seconds for 50-taxon networks, whereas the running time jumps to over 100 seconds for phylogenetic networks with 100 taxa. This increase in the running time with the number of taxa is expected, since the number of trinets of a network on *n* taxa is 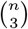. The very small increase in the running time, by viewing it as a function of the increase in the number of reticulations, is because trinets are small enough for full-likelihood calculations. Furthermore, even when a network has 4 reticulations, many trinets would have 0 or 1 reticulations, making their analysis even faster.

While scalability in speed is impressive, the main question is: How accurate is inference of phylogenetic networks under pseudo-likelihood when using bi-allelic markers? To answer this question, we generated 100 data sets, 20 replicates for each number of sites on the network of Fig. 4. We simulated 100, 1000, 10000, 100000, and 1000000 bi-allelic sites, with one haploid generated for every taxon. We set the mutation rates to *u* = 1 and *v* = 1 (where *u* and *v* are the mutation rate from red allele to green allele and the mutation rate from green allele to red allele, respectively, following the notation of [3]), then we used *θ* = 0.01 for every branch in the true network. We used following command (*numsites ∈ {*100, 1000, 10000, 100000, 1000000*}*) to simulate sites:

**Figure 4:**
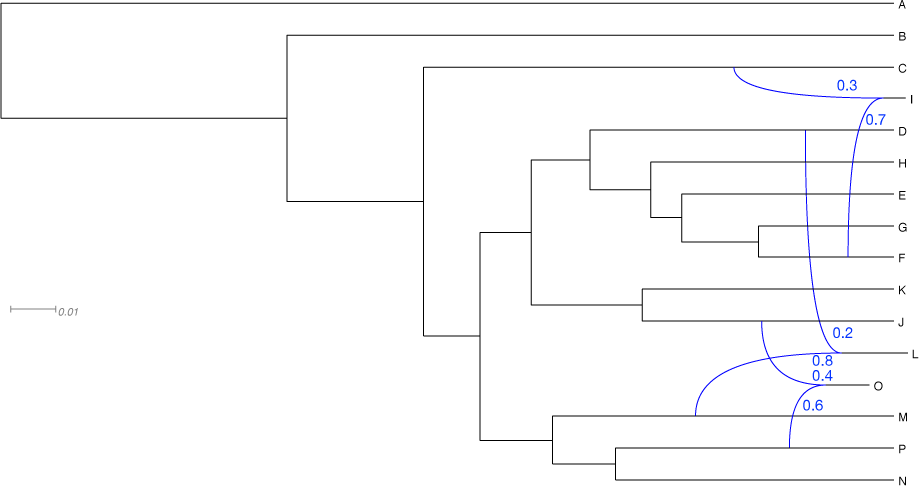
The true network with 16 taxa used for assessing the accuracy of inferences based on pseudo-likelihood. The branch lengths of the phylogenetic networks are measured in units of expected number of mutations per site (scale is shown). The inheritance probabilities are marked in blue.

~~~
SimBiMarkersinNetwork -pi0 0.5 -sd seed -num numsites -tm <A:A_0; B:B_0; C:C_0;D:D_0;E: E_0;F:F_0;G:G_0;H:H_0;I:I_0;J:J_0;K:K_0;L:L_0;M:M_0;N:N_0;O:O_0;P:P_0> -truenet
netstring -out "markers.txt";
~~~

For the value of *seed* in the “-sd” option, we used a different 8-digit integer for each of the 20 replicates. The value of *netstring* is the extended Newick string of the network in Fig. 4.

To test the ability of our algorithm to recover the topology of the true network, we ran the aforementioned simulated annealing procedure on the simulated data sets. The maximum number of reticulations during the search was set to 3, since determining the true number of reticulation is beyond the capability of a maximum unpenalized (pseudo-)likelihood [29]. For each data set, the search was performed for 50 iterations. The 5 networks with highest pseudo-likelihood were saved during the search. We used the following command to generate the results:

~~~
MLE_BiMarkers -pseudo -mnr 50 -pi0 0.5 -mr 3 -pl 8 -ptheta 0.01 -thetawindow 0.01 -sd 12345678 -taxa (A_0,B_0,C_0,D_0,E_0,F_0,G_0,H_0,I_0,J_0,K_0,L_0,M_0,N_0,O_0,P_0) -tm
<A:A_0; B:B_0; C:C_0;D:D_0;E:E_0;F:F_0;G:G_0;H:H_0;I:I_0;J:J_0;K:K_0;L:L_0;M:M_0;N: N_0;O:O_0;P:P_0>
~~~

The results are shown in Fig. 5. As the results show, when 10,000 sites or more are used as input for inference, the method always infers the true phylogenetic network. When only 1,000 sites are used, the true network is one of the top five optimal networks inferred, but not the most optimal, in 20% of the cases, whereas the true network is not even among the top five optimal networks in the remaining 80% of the cases. When only 100 sites are used, the method cannot infer the true network.

**Figure 5:**
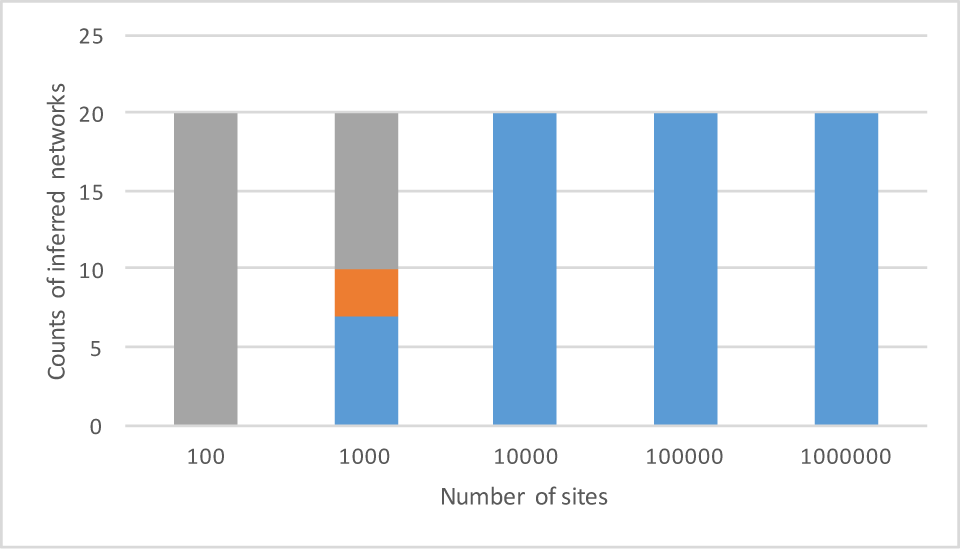
Accuracy of the inference on simulated data. The blue region corresponds to the number of times the true network was returned as the optimal network (the one with the highest pseudo-likelihood) after the search. The orange region corresponds to the number of times the true network is not the optimal network found by the search, but but is among the top 5 species networks under maximum pseudo-likelihood. All other scenarios are represented by the grey region.

These results can be put in the context of the performance of the method of [36] in terms of how inference based on the full-likelihood calculations performs, and the method of [31] in terms of how inference based on pseudo-likelihood from gene tree estimates performs. As shown by [36], accurate inference based on full-likelihood required at least 1,000 sites, even though the network considered in that work had only five taxa. In this regard, and given that the network considered here is much larger, it only makes sense that more than 1,000 sites are required. As for the method of [31], the authors showed that over 250 accurately estimated gene trees are required for accurate estimates of a 23-taxon phylogenetic network. An accurately estimated gene tree contains much more information than a single bi-allelic marker. In fact, a 23-taxon binary gene tree is equivalent to at least 21 different, yet compatible, bi-allelic markers, as each internal edge in the gene tree requires a bi-allelic marker to identify it.

These two contexts combined show the inference based on the pseudo-likelihood from bi-allelic markers is not only very accurate, but is also competitive with full-likelihood-based inferences. It is worth repeating that the method of [31] requires very accurate gene tree estimates. It is not uncommon to have error rates upwards of 60% in gene tree estimates on closely related species.

### 4.2. Comparison to full likelihood computation

To compare inferences based on pseudo-likelihood and full likelihood as given by the method of [36], we simulated 100, 1000, 10000, 100000, and 1000000 bi-allelic sites on the network in Fig. 6 with one haploid generated for every taxon. We used *θ* = 0.006 as the population mutation rate for external branches and the root, and *θ* = 0.005 for internal branches, both in the unit of population mutation rate per site. We set the mutation rates to *u* = 1 and *v* = 1. We generated 100 data sets, 20 replicates for each number of sites. We used following command (*numsites ∈ {*100, 1000, 10000, 100000, 1000000*}*) to simulate sites:

**Figure 6:**
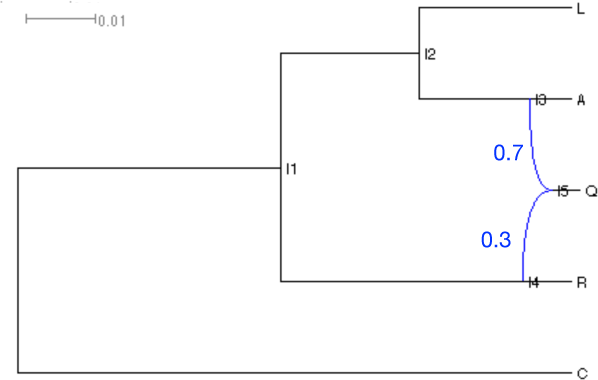
The true network with 5 taxa used for assessing the accuracy of inferences based on pseudo-likelihood. The branch lengths of the phylogenetic networks are measured in units of expected number of mutations per site (scale is shown). The inheritance probabilities are marked in blue.

~~~
SimBiMarkersinNetwork -pi0 0.5 -sd seed -num numsites -tm <A:A_0;C:C_0;L:L_0;Q:Q_0;R:R_0
> -truenet netstring -out "markers.txt";
~~~

For the value of *seed* in the “-sd” option, we used a different 8-digit integer for each of the 20 replicates. The value of *netstring* is the extended Newick string of network in Fig. 6.

Then we ran maximum likelihood inference on those data sets with 10 iterations with following commands, using both pseudo-likelihood and full likelihood computations (“-pseudo” is removed for the latter):

~~~
MLE_BiMarkers -pseudo -mnr 10 -pi0 0.5 -mr 1 -pl 8 -ptheta 0.006 -thetawindow 0.006 -sd 12345678 -taxa (A_0,C_0,L_0,R_0,Q_0) -tm <A:A_0; C:C_0;L:L_0;Q:Q_0;R:R_0>
~~~

Results of the comparison are shown in Fig. 7. As the results show, when 10,000 sites or more are used, inference based on pseudo-likelihood results in very accurate results that are identical to those obtained by full likelihood inference. When only 1000 sites were used, only in 6 out of 20 cases did pseudo-likelihood inference not result in the true network, but in all 6 cases that true network was one of the top 5 inferred. Again, in this case, the results are comparable to those based on full likelihood. In data sets consisting of only 100 sites, inferences result in poor networks regardless of whether full- or pseudo-likelihood is used. To summarize these results, inference based on pseudo-likelihood is comparable in accuracy to that based on full likelihood.

**Figure 7:**
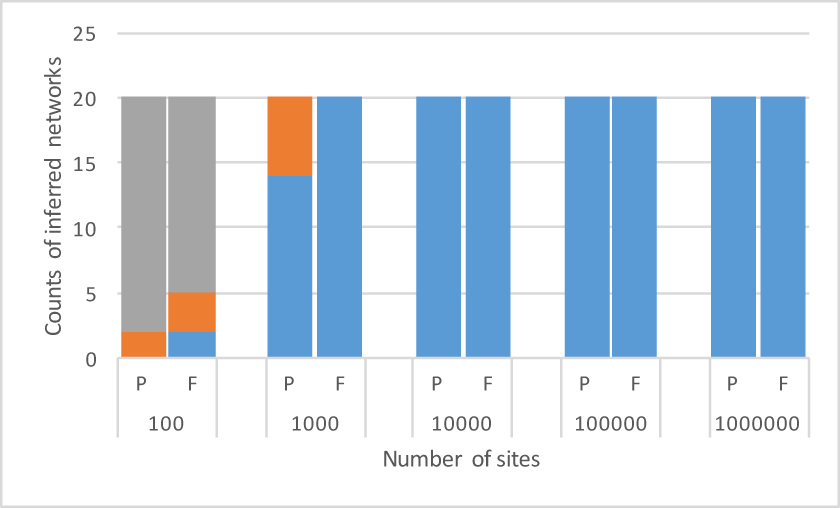
Accuracy of the inference on simulated data: pseudo-likelihood vs. full likelihood. The blue region corresponds to the number of times the true network was returned as the optimal network (the one with the highest pseudo-likelihood) after the search. The orange region corresponds to the number of times the true network is not the optimal network found by the search, but is among the top 5 species networks. All other scenarios are represented by the grey region. P: pseudo-likelihood, F: full likelihood.

### 4.3 The effect of number of individuals sampled

As described in [36], sampling more individuals from the hybrid species helps improve the accuracy of the inferred network based on full-likelihood computations. In particular, sampling more individuals allow the usage of fewer sites, a result that has important practical implications. We now set out to study this trend in the case of inferences based on pseudo-likelihood. We sampled one haploid individual for each of the four taxa L, A, R, and C, and 1, 2, 3, 4 haploid individuals for hybrid taxon Q in the network in Fig. 6. We generated 100, 500, 1000, 2000 sites for each of the 4 individual settings, with 10 replicates. We used following command (*numsites ∈ {*100, 500, 1000, 2000*}*) to simulate sites (in option “-tm”, individuals of Q: “Q 1,Q 2,Q 3” were removed according to 4 individual settings):

~~~
SimBiMarkersinNetwork -pi0 0.5 -sd seed -num numsites -tm <A:A_0; C:C_0;L:L_0;Q:Q_0,Q_1, Q_2,Q_3;R:R_0> -truenet netstring -out "markers.txt";
~~~

For the value of *seed* in the “-sd” option, we used a different 8-digit integer for each of the 20 replicates. The value of *netstring* is the extended Newick string of network in Fig. 6.

Then we ran maximum pseudo-likelihood estimation with 10 iterations on each of those 160 data sets using the same command as above. The accuracy of the inferred networks is reported in Fig. 8.

**Figure 8:**
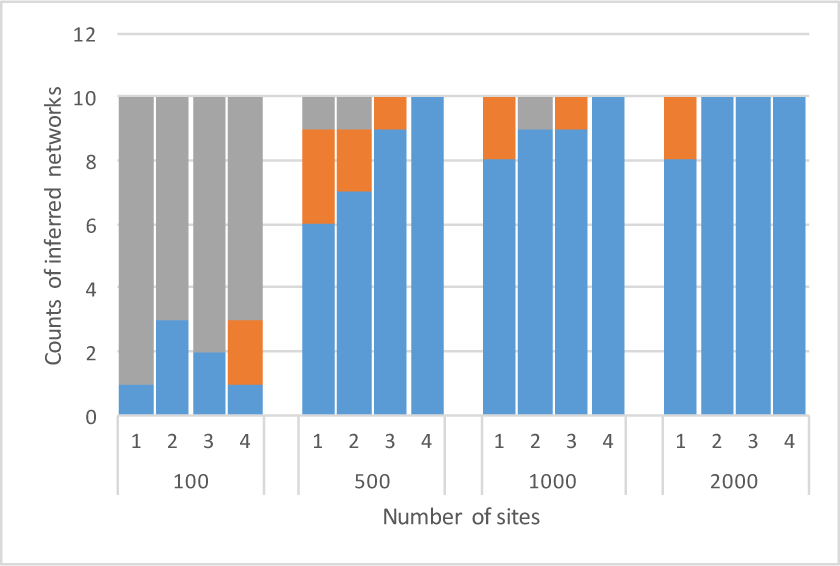
Accuracy of the inference on simulated data: Effect of the number of sampled individuals. The blue region corresponds to the number of times the true network was returned as the optimal network (the one with the highest pseudo-likelihood) after the search. The orange region corresponds to the number of times the true network is not the optimal network found by the search, but is among the top 5 species networks. All other scenarios are represented by the grey region. On the horizontal axis, 1, 2, 3 and 4 represent the number of individuals of sampled from the hybrid taxon.

The results show a clear benefit to sampling multiple individuals, especially when the number of sites is small. Once again, when only 100 sites are used, the performance is not very good, since this number of sites is too small. However, when only 500 sites are used, sampled only individual results in obtaining the true network in only 6 out of 10 cases, whereas sampling 4 individuals results in accurate inferences in all 10 cases. Furthermore, as the number of sites increases, the positive effect of increasing the number of sampled individuals starts diminishing.

### 4.4 Robustness of inference to lack of independence

As given by Eq. (1), the individual sites are assumed to be independent. Here, we set out to study the accuracy of inference when this assumption is violated. We simulated dependent sites as done by [36]. We generated 1000, 5000, and 10000 sites in the network of Fig. 6 under different conditions: (a) single marker was generated from a gene tree (this is the case of independent loci), (b) 10 markers were generated from a gene tree, (c) 100 markers were generated from a gene tree. Note that gene trees were generated independently. We used following command ((*numgt, sitespergt*) *∈ {* (1000, 1), (5000, 1), (10000, 1), (100, 10), (500, 10), (1000, 10), (10, 100), (50, 100), (100, 100)*}*) to simulate sites:

~~~
SimBiMarkersinNetwork -pi0 0.5 -sd seed -num numgt -sitespergt numsitespergt -tm <A:A_0; C:C_0;L:L_0;Q:Q_0;R:R_0> -truenet netstring -out "markers.txt";
~~~

For the value of *seed* in the “-sd” option, we used a different 8-digit integer for each of the 20 replicates. The value of *netstring* is the extended Newick string of the network in Fig. 6.

we ran maximum pseudo-likelihood estimation with 10 iterations on each of those data sets using the same command as above. Results are shown in Fig. 9. The results clearly show that the method is very robust to violation in the independent-loci assumption when 5000 sites or more are used. In the case of 1000 sites, the performance gets affected negatively only slightly. This result has great practical implications: In practice, when a large number of sites is sampled, one cannot ensure the sites are independent. The results show that even if independence is violated for a large number of sites, the method is robust in that case. This is consistent with what the authors observed with full likelihood computation in [36].

**Figure 9:**
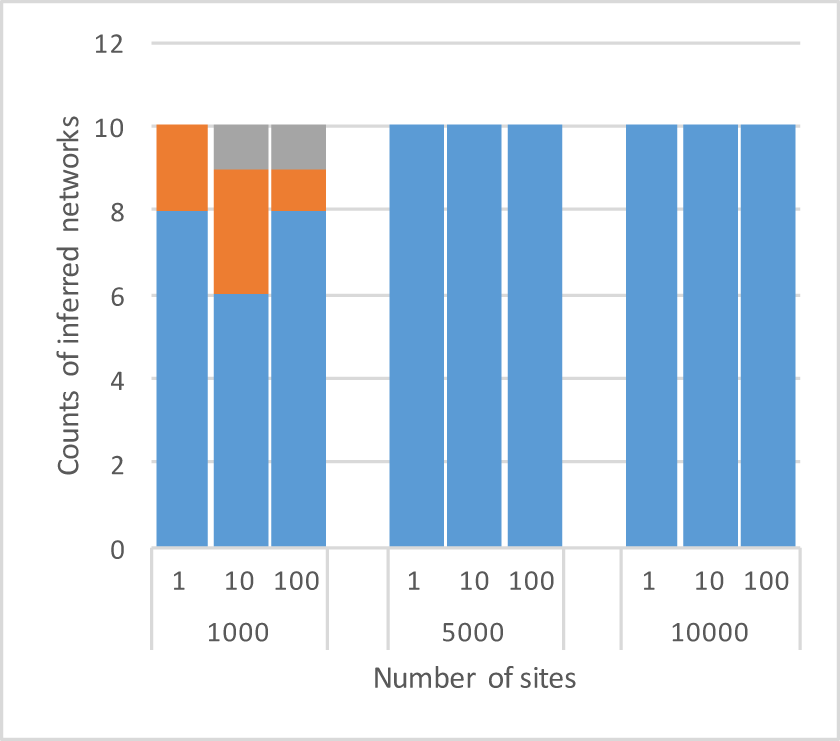
Accuracy of the inference on simulated data: Robustness to violation in the independent-loci assumption. The blue region corresponds to the number of times the true network was returned as the optimal network (the one with the highest pseudo-likelihood) after the search. The orange region corresponds to the number of times the true network is not the optimal network found by the search, but is among the top 5 species networks. All other scenarios are represented by the grey region. On the horizontal axis, 1, 10 and 100 represent the number of markers generated from a single gene tree.

### 4.5 The effect of limitation on number of reticulations

As we discussed in [29], inference of phylogenetic networks based on (unpenalized) likelihood cannot estimate the true number of reticulations simply because adding more reticulations only makes the model a better (or, at least as good a) fit for the data. This is why in the above results, we limited the number of reticulations that the method explores during inference to the true number (which is known since the data were simulated). To understand how the method performs when the limit on the number of reticulations during inference is set higher than the true number, we compared different limitations on the number of reticulations using the network in Fig. 6. We simulated 100, 1000, 10000, 100000, and 1000000 bi-allelic sites with 20 replicates for each number with the same command used for comparing pseudo-likelihood and full likelihood computation, then set the maximum number of reticulations to 1, 2 and 3 for each settings. We then ran maximum pseudo-likelihood inference with 10 iterations on those data sets using the same command as above, and the “-mr” (maximum reticulations) option is changed to 1, 2 and 3 accordingly. The results are shown in Fig. 10. The results make complete sense as setting a higher limit on the number of reticulations allowed during the search for optimal networks guides the method toward networks with more reticulations since those would have better pseudo-likelihoods. To illustrate the behavior of the method, we took a replicate with 10,000 sites as an example and plotted the pseudo-likelihoods of the best inferred networks with 1, 2, and 3 reticulations, as well as the networks themselves in Fig. 11. The figure shows two important points. First, as more reticulations are allowed, the pseudo-likelihood of the best networks found improves, but the improvement starts slowing down with the addition of more reticulations. Second, while more reticulations that the true number (one) are added, the true network is “inside” the optimal ones being identified with more reticulations. This is an important result because it illustrates how the true network is first identified and, then, when the method starts adding more reticulations, it does so to the underlying true network.

**Figure 10:**
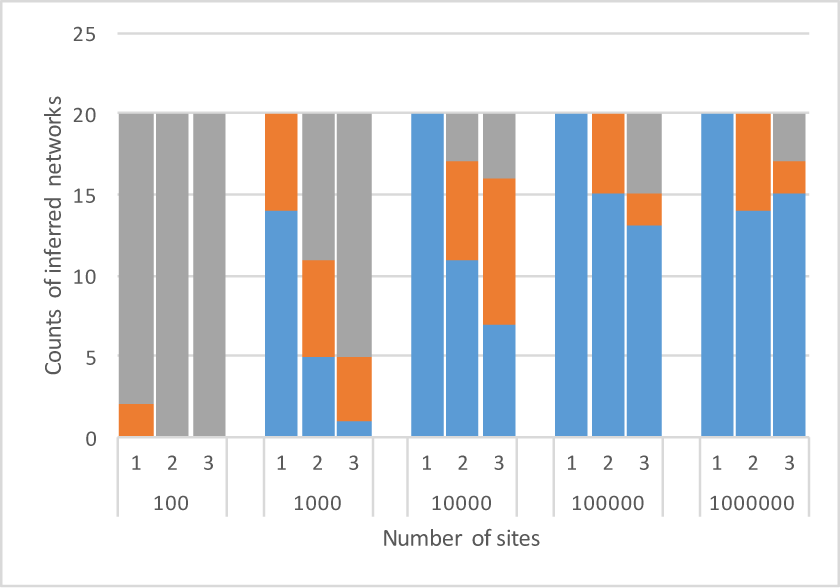
Accuracy of the inference on simulated data: Setting different limits on the number of reticulations during inference. The blue region corresponds to the number of times the true network was returned as the optimal network (the one with the highest pseudo-likelihood) after the search. The orange region corresponds to the number of times the true network is not the optimal network found by the search, but is among the top 5 species networks. All other scenarios are represented by the grey region. On the horizontal axis, 1, 2 and 3 represent the maximum number of reticulations during inference.

**Figure 11:**
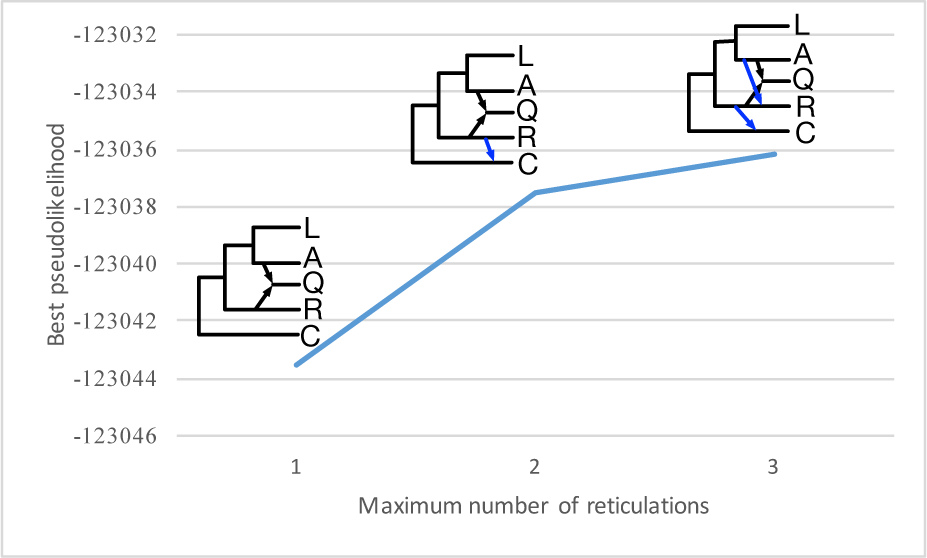
Improvement with more reticulations. On the horizontal axis, 1, 2 and 3 represent the maximum number of reticulations during the estimation. At each point, the corresponding network topology is shown. The blue edges represent the additional reticulations to the true network.

### 4.6 Empirical data sets

We also analyzed the two data sets in [36] using our algorithm, including two different hybrid individuals *O. × cockayneana* and *O. × prorepens*. The hybrid origins of these two individuals are supported by both morphological [14] and molecular (Meudt unpubl.) data. These two individuals were formally named along with putative parents. The first data subset comprises the following five individuals: *O. macrocarpa* (voucher: *Meudt 133a*, MPN 29546; herbarium codes follow [25] [continuously updated]), *O. macrophylla* subsp. *lactea* (*Cameron 13392*, AK 294893), hybrid *O. × cockayneana* (*Meudt 175a*, MPN 29710), *O. caespitosa* (*Meudt 174a*, MPN 29705), and *O. calycina* (*Meudt 176a*, MPN 29713). The number of loci in this data set is 802. The second data subset comprises *O. sessilifolia* subsp. *splendida* (*Heenan s.n.*, MPN 32149), *O. macrocarpa* (*Meudt 133a*, MPN 29713), hybrid *O. × prorepens* (*Meudt 203<u>^a^</u>*, MPN 29774), *O. sessilifolia* subsp. sessilifolia (*Meudt 199a*, MPN 29771), and *O. caespitosa* (*Meudt 196a*, MPN 297695). The number of loci in this data set is 820.

Each data subset comprised five diploid individuals in total, which means ten haploid individuals were effectively analyzed due to the correction for dominant markers. Observe that while we could combine the two data sets into a larger one to study the scalability of the method on a larger data set, the number of sites available is too small for accurate inferences by pseudo-likelihood, as supported above by the simulation study.

The search was performed for 50 iterations for each data set. The maximum number of reticulations was set to 1. The results are given in Fig. 12 and Fig. 13.

**Figure 12:**
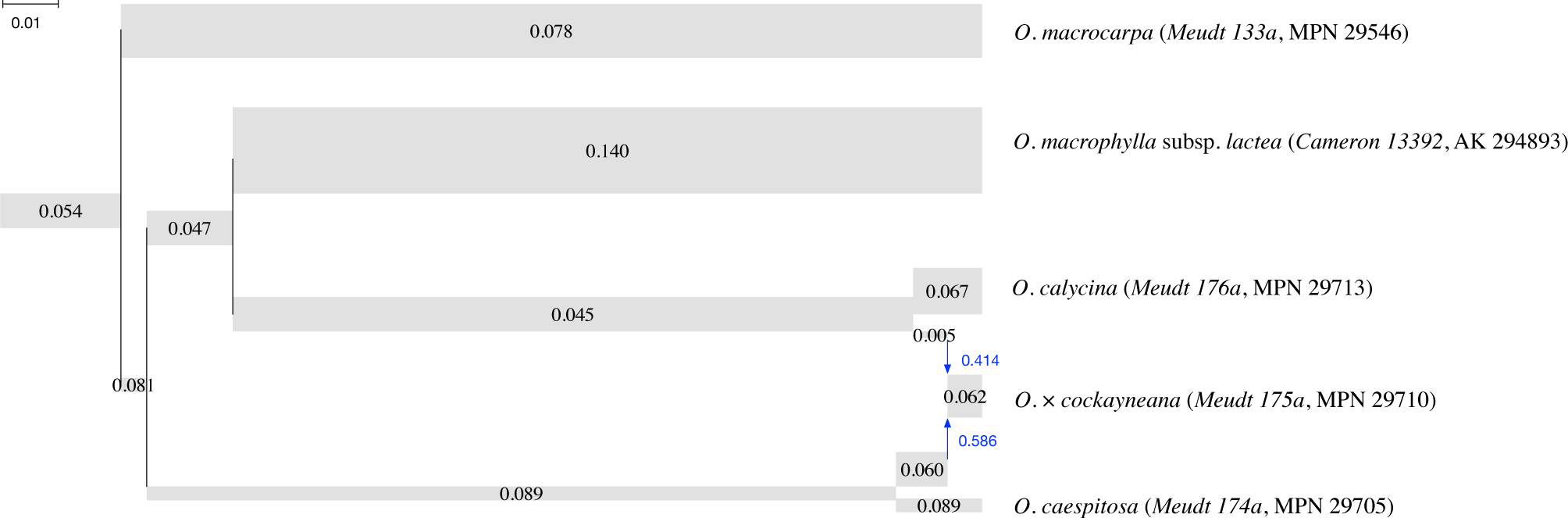
The phylogenetic network with maximum pseudo-likelihood for the subset with the hybrid *O. × cockayneana* (*Meudt 175a*, MPN 29710) and putative parents. The width of each tube is proportional to the population mutation rate of each branch, which is printed on each tube. The length of each tube is proportional to the length of the corresponding branch in units of expected number of mutations per site (scale shown). Blue arrows indicate the reticulation edges and their inheritance probabilities are printed in blue.

**Figure 13:**
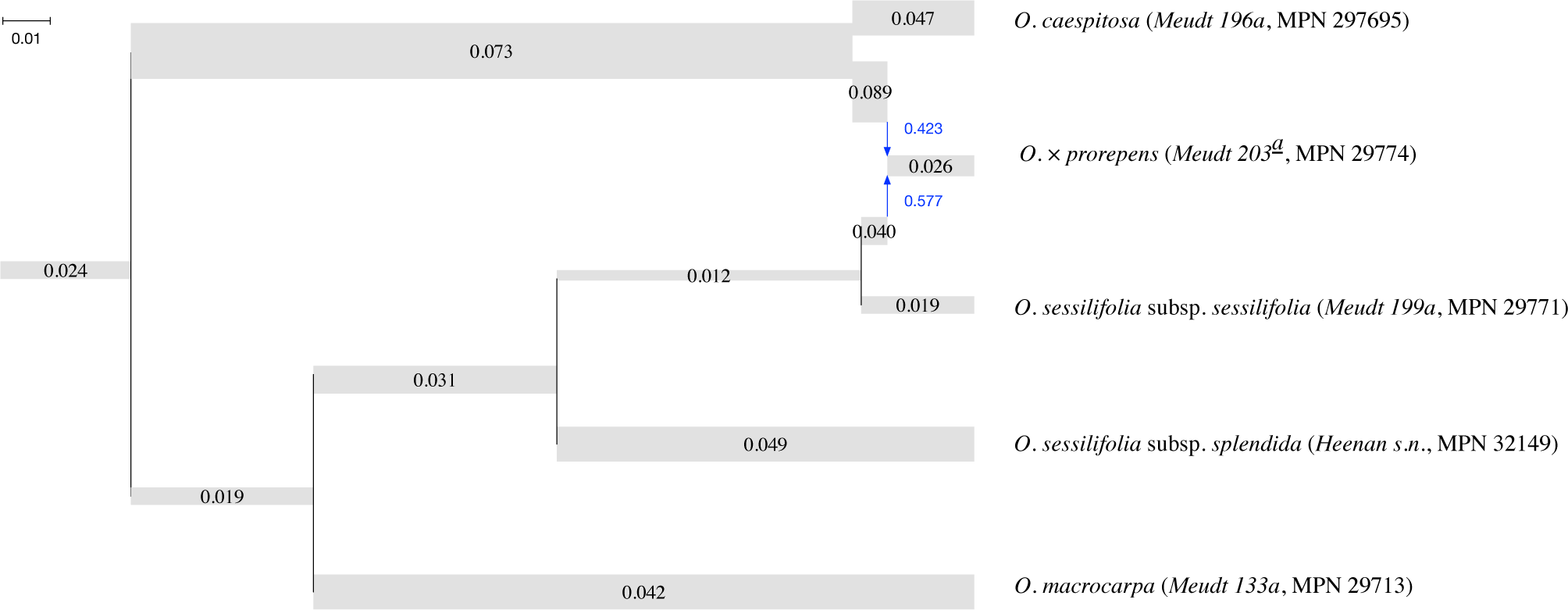
The phylogenetic network with maximum pseudo-likelihood for the subset with the hybrid *O. × prorepens* (*Meudt 203*<u>^*a*^</u>, MPN 29774) and putative parents. The width of each tube is proportional to the population mutation rate of each branch, which is printed on each tube. The length of each tube is proportional to the length of the corresponding branch in units of expected number of mutations per site (scale shown). Blue arrows indicate the reticulation edges and their inheritance probabilities are printed in blue.

Both results show that hybrids are correctly detected. The topology of the network in Fig. 13 with the hybrid *O. × prorepens* is consistent with Fig. 20 in [36], while the topology of the network in Fig. 12 with the hybrid *O. × cockayneana* is slightly different from that in Fig. 19 in [36]. Notably, the topology of Fig. 19 in [36] is inferred as the second-best network by our algorithm, and its pseudo-likelihood value is smaller than that of the best network by only 0.2.

## 5 Discussion

The results above on simulated data and empirical data demonstrate very good accuracy of phylogenetic network inference using pseudo-likelihood based on bi-allelic markers. In terms of computing the pseudo-likelihood, the computation scales up to hundreds of taxa. Of course, searching the space of 100-taxon networks is a different challenge that requires novel techniques beyond scaling up likelihood computations. The simulation results, in particular, clearly demonstrate that as the number of bi-allelic markers increases, the accuracy improves significantly. We now turn to investigating the convergence of the inferred phylogenetic network onto the true network as the number of sites increases.

Let Ψ be a phylogenetic network on set 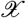 of taxa and consider a subset 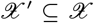 with 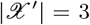. Both Ψ and trinet 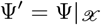 can be viewed as generative models for bi-allelic markers. It is important here to remind the reader that all parameters of Ψ′are transferred from Ψ. Let *g* be a gene tree on set 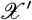 of taxa. We have *p*(*g|*Ψ) = *p*(*g|*Ψ′), since the density *p*(*g|*Ψ) does not involve any branches (or their parameters) that are in Ψ but not in Ψ and the length of a branch in Ψ corresponds to the length of either a branch in Ψ or a path that results from removing nodes when obtaining Ψ′. It follows from this that *P* (*s|g*)*p*(*g|*Ψ) = *P* (*s|g*)*p*(*g|*Ψ′) for a bi-allelic marker on the set 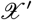 of taxa. Therefore, we obtain from this that *P* (*s|*Ψ) = *P* (*s|*Ψ′). In other words, as the number of sites goes to infinity, for every trinet of the true network, the proportions of marginalized site patterns converge to their expectation in this trinet of the true network.

To study the convergence empirically, we generated one data set with each of 100, 1000, 10000, 100000, 1000000 sites of bi-allelic markers using the network of Fig. 4. Then for each data set, for every trinet of that network, we compute the proportion of every corresponding marginalized site pattern, and compare it to the expected frequency of that pattern in the trinet. We plotted the differences between these proportions in Fig. 14.

**Figure 14:**
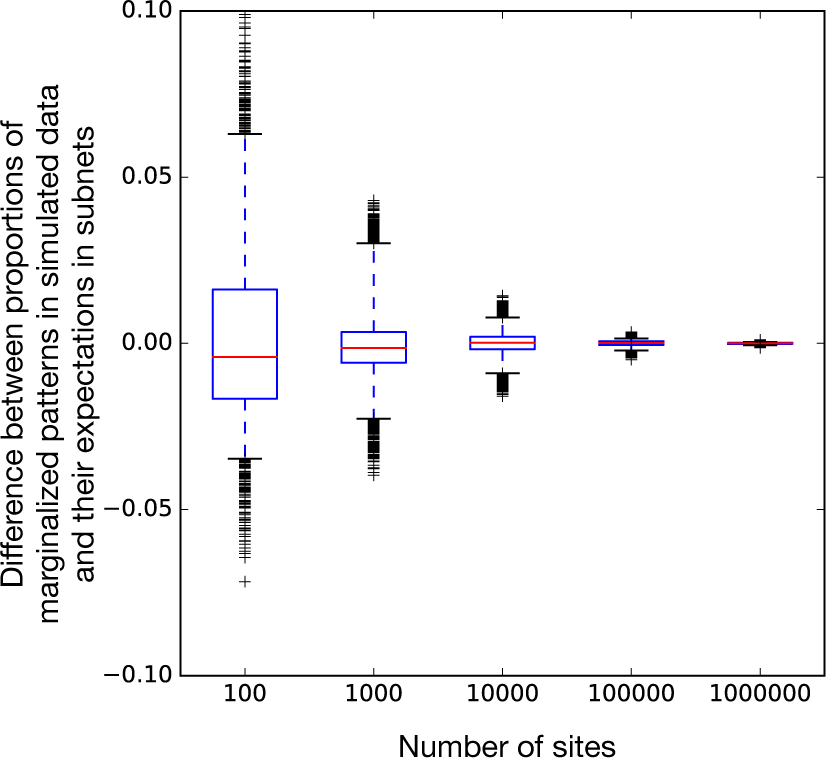
Convergence of the proportions of marginalized site patterns in the data to their expectations. Every point is the empirical frequency of a marginalized 3-taxon site pattern minus the (theoretical) expectation of that frequency on the corresponding subnet in the true network.

As the results show, the proportions of site patterns of trinets converge very fast to their theoretical expectation, with very negligible variance in the differences once 100,000 sites or more are used. These results are compatible with the accuracy of the method discussed above on the simulated data.

## 6 Conclusions

Statistical approaches for inferring phylogenetic networks offer a great promise in terms of utilizing the data to estimate not only the topology of the network, but also evolutionary parameters of interest to the biologist, such as population mutation rates and divergence times. Statistical approaches are based on evaluating the likelihood of phylogenetic network candidates during search of the parameter space. Except for small networks with fewer than ten taxa and two or three reticulations, these approaches are hard to apply in practice given the prohibitive computational requirements. The pseudo-likelihood methods of [31] and [21] offered a way to ameliorate this issue. However, these methods make use of gene tree estimates as the input data. Under conditions of extensive incomplete lineage sorting, the individual loci could have very little signal to obtain accurate gene trees. The major contribution of this paper is the introduction of a pseudo-likelihood method based on the sequence data directly. We demonstrated that evaluating the pseudo-likelihood of a phylogenetic network is very fast and allows scaling to very large networks. We also demonstrated the accuracy of maximum pseudo-likelihood inference of phylogenetic networks from bi-allelic data.

Even though the pseudo-likelihood formulation provides accurate inferences, it still does not circumvent the challenging problem of searching the space of large networks. Developing more efficient moves for walking the space of phylogenetic networks is necessary.

## Acknowledgements

The authors would like to thank Heidi Meudt for sharing data for the two biological data sets.

